# NEPRE: a Scoring Function for Protein Structures based on Neighbourhood Preference

**DOI:** 10.1101/463554

**Authors:** Siyuan Liu, Xilun Xiang, Haiguang Liu

## Abstract

Protein structure prediction relies on two major components, a method to generate good models that are close to the native structure and a scoring function that can select the good models. Based on the statistics from known structures in the protein data bank, a statistical energy function is derived to reflect the amino acid neighbourhood preferences. The neighbourhood of one amino acid is defined by its contacting residues, and the energy function is determined by the neighbhoring residue types and relative positions. A scoring algorithm, Nepre, has been implemented and its performance was tested with several decoy sets. The results show that the Nepre program can be applied in model ranking to improve the success rate in structure predictions.

## INTRODUCTION

Despite the advances in protein structure determination methods, the discovery rate of new proteins far exceeds the speed of experimental structure determination for protein molecules. New proteins can be automatically discovered from high throughput genome sequencing data with sophisticated genome analysis tools (Bateman *et al.*, 2015; Kim *et al.*, 2014; Altelaar *et al.*, 2013). On the other hand, the protein structure determination requires complicated procedures to get high quality protein samples for good experimental signals. For example, the targeting protein must have a reasonable expression rate to obtain sufficient amount of samples, which are subsequently purified, followed by the optimization of crystallization cocktail recipes to yield high quality crystals in the case of X-ray crystallography (Carpenter *et al.*, 2008; Slabinski *et al.*, 2007), or molecules have to be labelled using isotopes at specific atoms in the case of nuclear magnetic resonance (Markwick *et al.*, 2008; Billeter *et al.*, 2008). The recent breakthrough in cryogenic electron microscopy methods shines light on the possibility of high throughput structure determination (Cheng, 2018). However, the technology is still not highly automated and requires extensive computational analysis on large volume of data. The limitation on experimental structure determination of protein molecules urges the development of protein structure prediction using computational modelling method.

The protein structure prediction research has a long history, marked by the popular international prediction contests, the Critical Assessment of protein Structure Prediction, the CASP that was first organized in 1994 (Moult, 2005; Moult *et al.*, 2018). The structure prediction has achieved successes in many cases and attracted numerous applications (Zhang, 2008). In particular, the predicted structures can be combined with experimental data to provide more comprehensive understanding of the molecules (Nealon *et al.*, 2017; Schneidman-Duhovny *et al.*, 2012; Dos Reis *et al.*, 2011; Latek *et al.*, 2007; Wang and Liu, 2017). In many cases, it is difficult to determine a high quality structure with limited experimental information alone. The hybrid methods that integrate structure prediction results and experimental data are very promising to exploit the information from both experimental research and computational predictions. For a structure prediction method to be successful, it must have two components: (1) an algorithm to generate an structure ensemble that include good models, i.e., at least some models in the ensemble are similar to the correct structure; and (2) a scoring function that can rank the generated structures, so that the good models can stand out from the rest. The structure ensemble for a protein, often referred to as a decoy set, can be generated using several computational methods. The main stream methods include homology modeling (Martí-Renom *et al.*, 2000), structure threading (Lemer *et al.*, 1995; Xu *et al.*, 2010), and segments assembling (Rohl *et al.*, 2004; Lange and Baker, 2012; Lee *et al.*, 2011). The advanced sampling algorithms can be applied to ensure diversity of the decoy set, such that the chance of sampling the best structure can be guaranteed (Lee *et al.*, 2011). In this work, we focus on the scoring function that used to assess the quality and correctness of each generated model.

There are two types of scoring functions, one is based on physical chemistry principles, represented with the force fields in molecular modeling, such as Amber or Charmm for atomic models (Case *et al.*, 2005; Brooks *et al.*, 2009) and Martini or UNIRES for coarse grained models (Marrink *et al.*, 2007; Monticelli *et al.*, 2008; Liwo *et al.*, 1999). The other type is empirical energy functions based on statistics from knowledge of experimentally determined structures. There has been tremendous success in applying these empirical energy functions to predict protein structures. One famous example is the protein main chain dihedral angle distributions, known as Ramachandran plot (Ramachandran and Sasisekharan, 1968), which is widely used for protein structure validation (Laskowski *et al.*, 1993; Hooft *et al.*, 1997; Davis *et al.*, 2004). The outstanding developments in empirical energy functions include DFIRE, DOPE, RW, RWplus, GOAP, etc (Zhang, 2004; Shen and Sali, 2006; Zhang and Zhang, 2010; Zhou and Skolnick, 2011). Inspired by these pioneer work, we developed a new energy function that describes the amino acid neighborhood preferences. For each of the 20 natural amino acids, the neighboring amino acid was analyzed in detail. Specifically, the preference was understood using 400 (20×20) matrices that describe the relative positioning of any two types of amino acids. The likelihood to be neighbors for different types of amino acids was also counted in the implementation. First, for any two types of amino acids, the likelihood to be neighbors and the likelihood to be neighboring in each discretized section in spherical coordinate were extracted from a high resolution structure dataset. The likelihood distributions were converted to energy functions using Boltzmann relation, and these energy functions were used to evaluate the decoy structures. Based on the testing results and the comparison with several other structure assessment methods, we report that the neighborhood preference (Nepre) function is effective in ranking the decoy structures and quantifying the structural correctness.

## MATERIALS AND METHODS

The native state structures of proteins are mostly stabilized by the weak interactions between atoms that are not covalently bonded, mostly include electrostatic or van der waal interactions. Although these weak interactions are nonspecific, each residue is found to have preferences on its neighboring residue types, especially on the nearest neighbors. Furthermore, the relative position of the neighboring residues are also critical for their packing. With this in mind, we carried out detailed statistics on the neighborhood preference of each amino acid type. First, a local coordinate system was defined for each amino acid to describe its neighboring residue positions; secondly, the neighboring residues were mapped to the polar coordinates defined around the center residue; at last, every amino acid in the protein molecule was treated as the center residue in turn to obtain the statistics of overall neighborhood preference. The final statistics were obtained from a non-redundant dataset composed of 14,647 PDB structures, which were selected from the NCBI VAST (the vector alignment search tool) server with p-value=10^−7^ (Gibrat *et al.*, 1996).

### Local coordinate system for each amino acid

The local coordinate system was defined using main chain atoms of each amino acid, as described in an earlier work (Xiang and Liu, 2018). In brief, the X-Y plane was defined using the geometry center (g_c_) of the focusing amino acid, nitrogen atom (N), and carboxyl carbon atom (C). The geometry center, g_c_, only accounts for the non-hydrogen atoms, and is defined as the origin point (O) of the local coordinate system. The positive x-direction is defined as O→N, then the positive y-direction can be defined in the X-Y plane such that the carboxyl atom has a positive y coordinate. The z-direction was defined using the right-hand rule.

The neighboring amino acids were defined based on the distances between the centers of the corresponding amino acid side chains. If the distance is within a given cutoff value, they are considered as the neighbors. Once the neighborhood is defined, the statistics is carried out within the cutoff, therefore, the scoring function becomes distance-independent at this level. We used two approaches for the distance cutoff: a universal fixed cutoff for all amino acids and the other one depends on the neighboring amino acid types.

For the case of fixed cutoff, the performance of the algorithm was tested using different cutoff values, with r_c_ between 4Å and 10Å. For the second case that the cutoff was determined by the radii of two neighboring amino acids. The radius distributions of 20 amino acids were studied from the same VAST structure dataset.

### The statistical model for amino acid contacts in protein molecules

The distribution function is related to the energy via the Boltzmann’s law, explicitly, the energy can be expressed as:

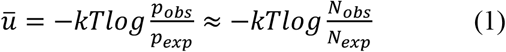

where the *p_obs_* and *p_exp_* are the observed and expected probabilities in the subspace specified with parameters of interest. In Nepre, *p_obs_* and *p_exp_* are defined with five parameters: (*i*, *j, r, θ, φ*). While (*i*, *j*) are the types of amino acids, (*r, θ, φ*) represent the relative coordinate parameters of the latter in the former amino acid’s spherical coordinate. To simplify the representations, the geometric center of each amino acid is used to represent its location in the centerred amino acid (See Figure 2). From the structure database, the observation of amino acid type *j* in the neighbourhood of amino acid *i* is expressed as *p_obs_*(*i*, *j, r, θ, φ*).

**Figure 1.**
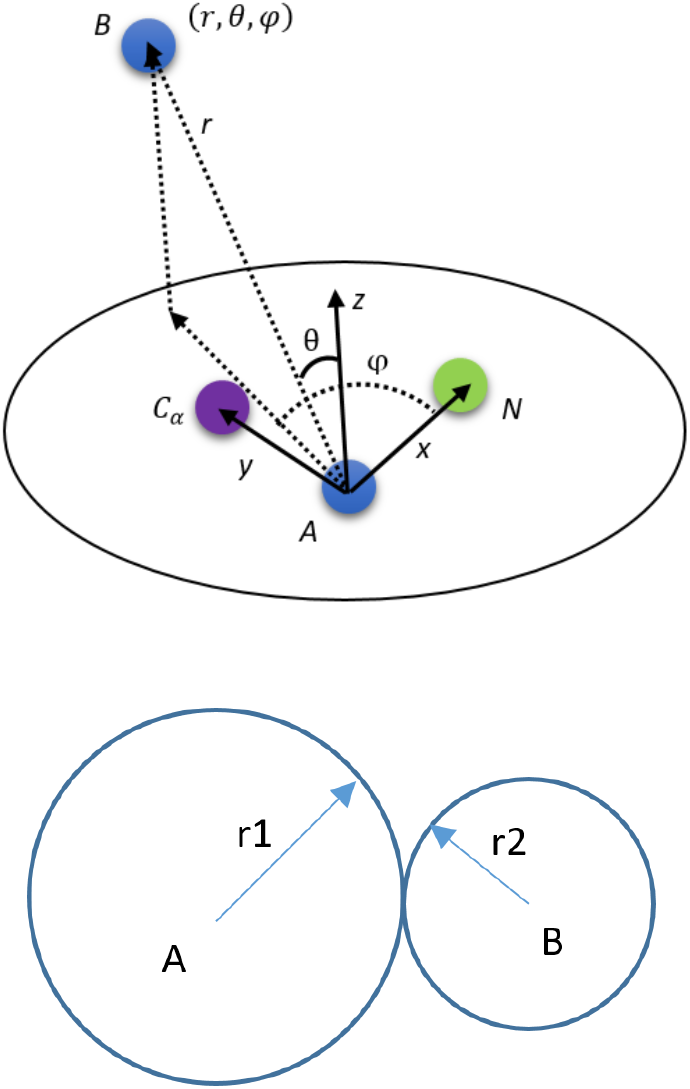
The schematic drawing of two neighboring amino acids. **(a)**The coordinate of amino acid B in coordinate system of amino acid A. (**b**) The distance between two amino acids.

**Figure 2.**
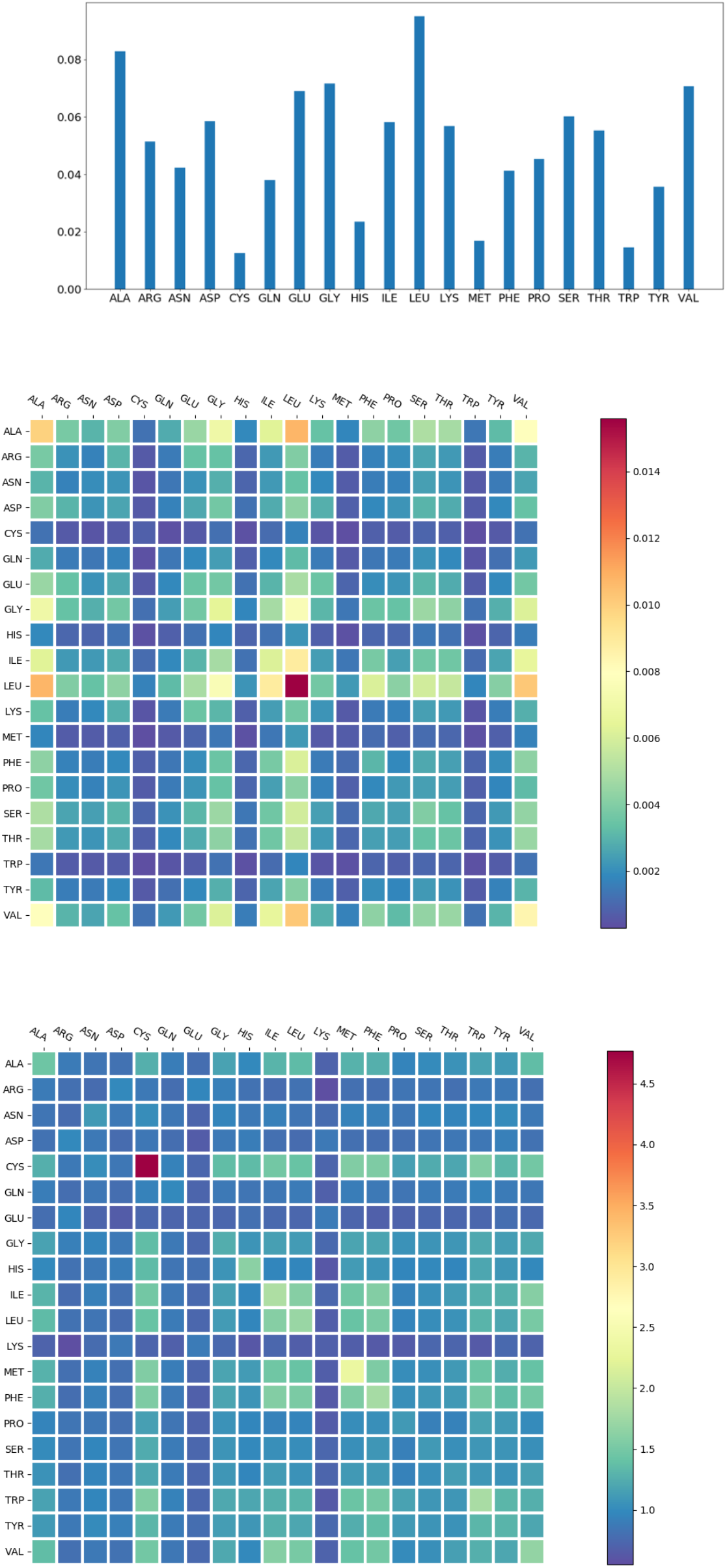
Probability of observing amino acid and amino acid neighbors. (**a**) amino acid abundance (normalized) in the protein dataset; (**b**) the probability of amino acid types that are neighboring in pairs; (**c**) the observation to expectation ratio for types of neighboring amino acids.

The observed probability *p_obs_*(*i*, *j, r, θ, φ*) can be expressed as:

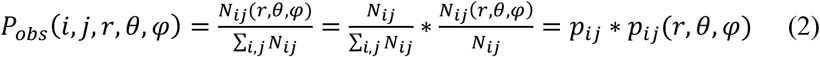

The expected values of the distribution of various amino acids are uniform, which can be expressed as:

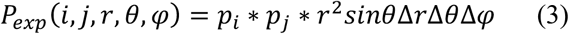

According to the above derivation, we got:

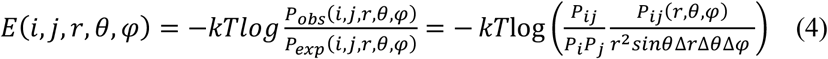

Where *k* is the Boltzmann constant and *T* is the temperature, (*r*, *θ*, *φ*) is the spherical coordinate of the amino acid *j* around the amino acid *i*.

For a protein with M amino acids, the energy can be expressed as:

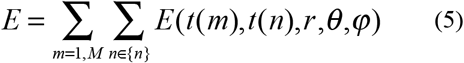

Where m is the index of the amino acid, {n} is the neighboring amino acid with the given cutoff value; t(x) is the function that maps the amino acid type to each amino acid.

In the implementation of the program, the distance was integrated out as the statistics were carried out in the sections specified by the angles (θ, *φ*) within the contacting sphere. A regular grid system was used to divide the sphere into 20×20 regions, with 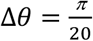 and 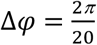 (because the range for θ is [0, *π*) and for φ is [0,2*π*)). Although this setup does not give an equal volume division, the effect can be corrected by using the appropriate probability in the respected volume (see equation 3).

### Testing decoy datasets

The performance of the algorithms were tested using five published datasets: the original I-Tasser dataset, denoted as I-Tasser(a), and four datasets generated using the 3DRobot programs, including I-Tasser(b), 3DRobot, Rosseta, and Modeller. The information about the datasets is summarized in Table 1.

**Table 1.**
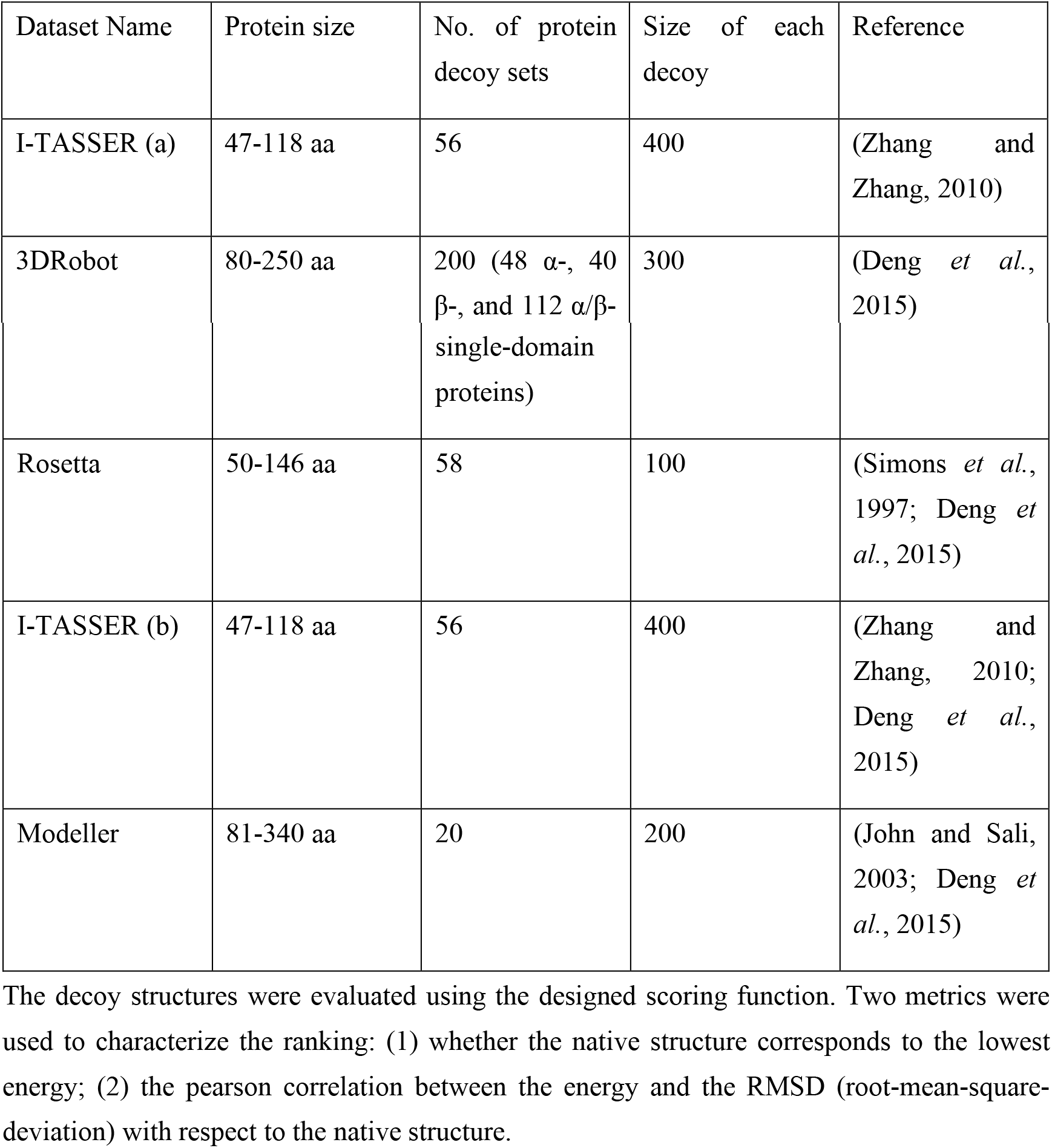
Summary of the five decoy datasets.

## Results

### The probability of neighboring for amino acids

The 20 natural amino acids appear in protein molecules with different abundances. The probability of finding a particular type of amino acid in the nonredundant dataset is summarized in Figure 2a, showing that the hydrophobic amino acids, such as leucine, alanine and valine, appear in protein molecules more frequently than others. The probabilities of observing two neighboring amino acids for the 20×20 pairs were also studied. Figure 2b shows the probability distribution for the cases with the distance cutoff value 6.0 Å. The neighborhood preference is quantified using the observed to expected ratio, o/e, defined as 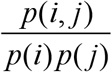, which is summarized in Figure 2c. It is clear that some amino acid types have strong preferences about their neighbors, such as cysteine strongly prefers another cysteine in its neighborhood. This information is useful to quantify the packing of amino acids in protein structures, with the applications explained in the method section.

### Location preferences of amino acids within the neighborhood

The amino acids interact with each other in their preferred orientations, as revealed by the uneven distribution of one amino acid within the sphere centered at another amino acid. This provides additional information on top of the preferred pairing discussed in the previous section. In figure 3a, the distribution of glutamate acid around proline is shown in a contour map, indicating that the proline most likely to be in the region around (63°,144°). Cysteines have even stronger preferences upon their cysteine neighbors, concentrated in the region of (162°, 270°) (Figure 3b).

**Figure 3.**
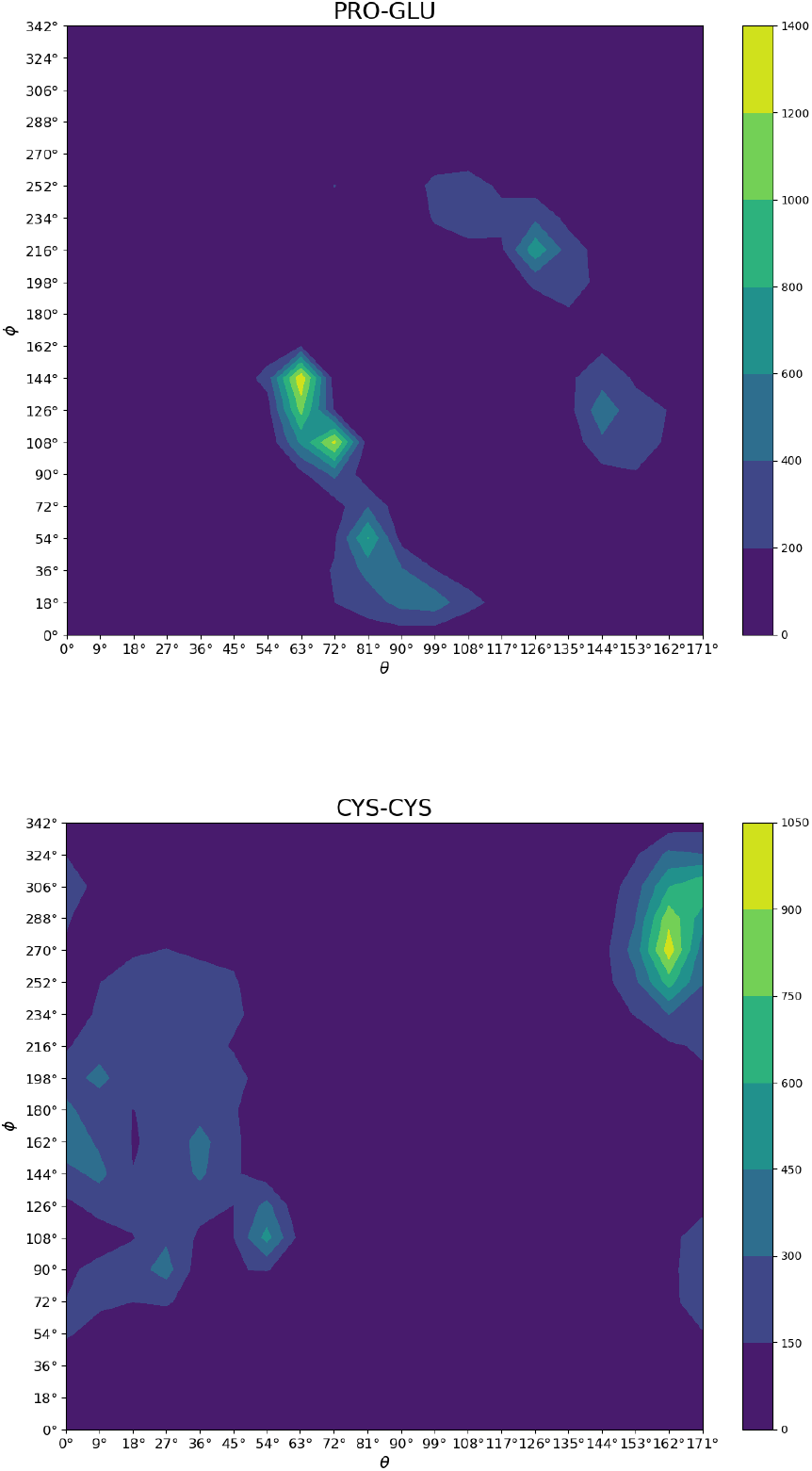
The distribution of amino acid in the neighborhood of each other. (**a**) glutamate acid around proline within the neighborhood of 6Å; (**b**) cysteine in the neighborhood of another cysteine.

### The performance of Nepre

As described in the Method section, the neighbourhood preference based scoring function (Nepre) has two implementations depending on the choice of neighbourhood cutoff values. One implementation utilizes a fixed cutoff value for all amino acid types, hereafter named as Nepre-F; and the second implementation has cutoff values depending on the neighboring amino acid types, and this one is named to be Nepre-R (meaning that the cutoff value depends on the radii of the neighboring amino acids).

For the case of Nepre-F, the cutoff value is critical for the neighborhood boundary. We tested the scoring function at various cutoff values from 4Å to 10Å for five datasets described in the Method section. The success rates for picking out the native states for each decoy set were summarized in Table 2.

**Table 2.** The number of success cases with different fixed cutoffs.

| Cutoff | I-TASSER(a) | 3DRobot | I-TASSER(b) | ROSETTA | Modeller |
| --- | --- | --- | --- | --- | --- |
| 4 Å | 11/56 | 27/200 | 4/56 | 8/58 | 9/20 |
| 5 Å | 42/56 | 95/200 | 16/56 | 27/58 | 12/20 |
| **6 Å** | **48/56** | **106/200** | **21/56** | **32/58** | **13/20** |
| 7 Å | 49/56 | 89/200 | 19/56 | 25/58 | 14/20 |
| 8 Å | 49/56 | 80/200 | 18/56 | 21/58 | 12/20 |
| 9 Å | 49/56 | 65/200 | 18/56 | 13/58 | 11/20 |
| 10 Å | 50/56 | 60/200 | 10/56 | 10/58 | 11/20 |

**Figure 4.**
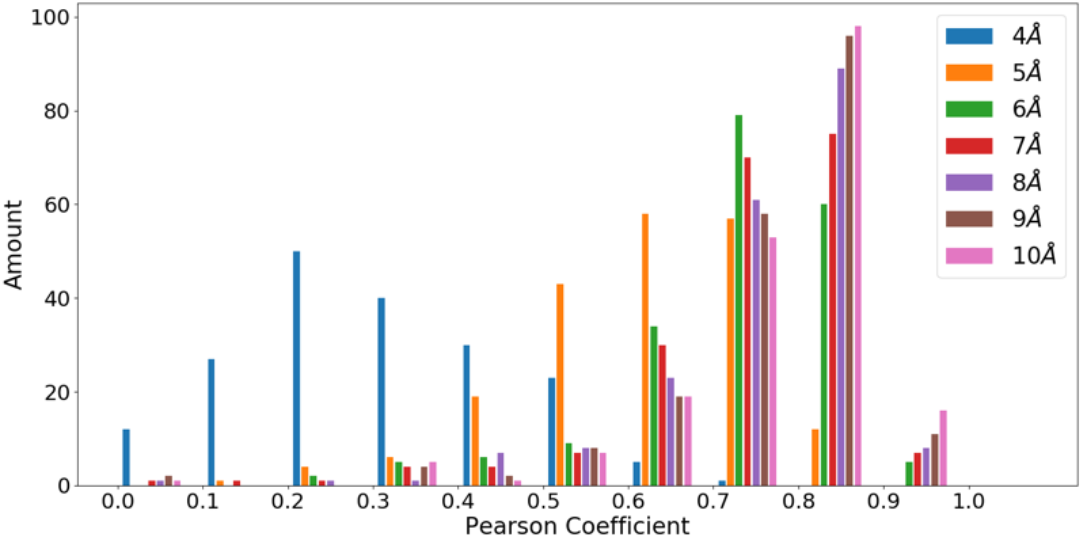
Overall ranking quality measured using pearson correlation coefficients. The distributions of pearson correlation coefficients are shown at cutoff values from 4Å to 10Å.

For the case of Nepre-R, the radii for each type of amino acid were extracted from the non-redundant dataset. The distributions of radii for 20 amino acids were shown in the supplementary figure S1 and the mean values were summarized in Table 3. These values were used to determine the cutoff values in the neighborhood statistics for specific amino acid types. From the neighborhood analysis, the associated energy function was derived as in the case of Nepre-F.

**Table 3.** The mean value of radius for each amino acid type.

| Amino Acid Type | Radius(Å) | Amino Acid Type | Radius(Å) |
| --- | --- | --- | --- |
| ALA | 3.20 | LEU | 4.24 |
| ARG | 5.60 | LYS | 5.02 |
| ASN | 4.04 | MET | 4.47 |
| ASP | 4.04 | PHE | 4.99 |
| CYS | 3.65 | PRO | 3.61 |
| GLN | 4.64 | SER | 3.39 |
| GLU | 4.63 | THR | 3.56 |
| GLY | 1.72 | TRP | 5.38 |
| HIS | 4.73 | TYR | 5.36 |
| ILE | 3.94 | VAL | 3.55 |

### Comparison of native structure selection using different potentials

In Table 4, we present the recall rates and the z-scores of Nepre-R and Nepre-F on the five decoy datasets. Overall, Nepre-R performance is better on I-TASSER(a) and I-TASSER(b) datasets with lower z-scores, while Nepre-F performs better on modeller, Rosetta, and 3DRobot datasets.

**Table 4.**
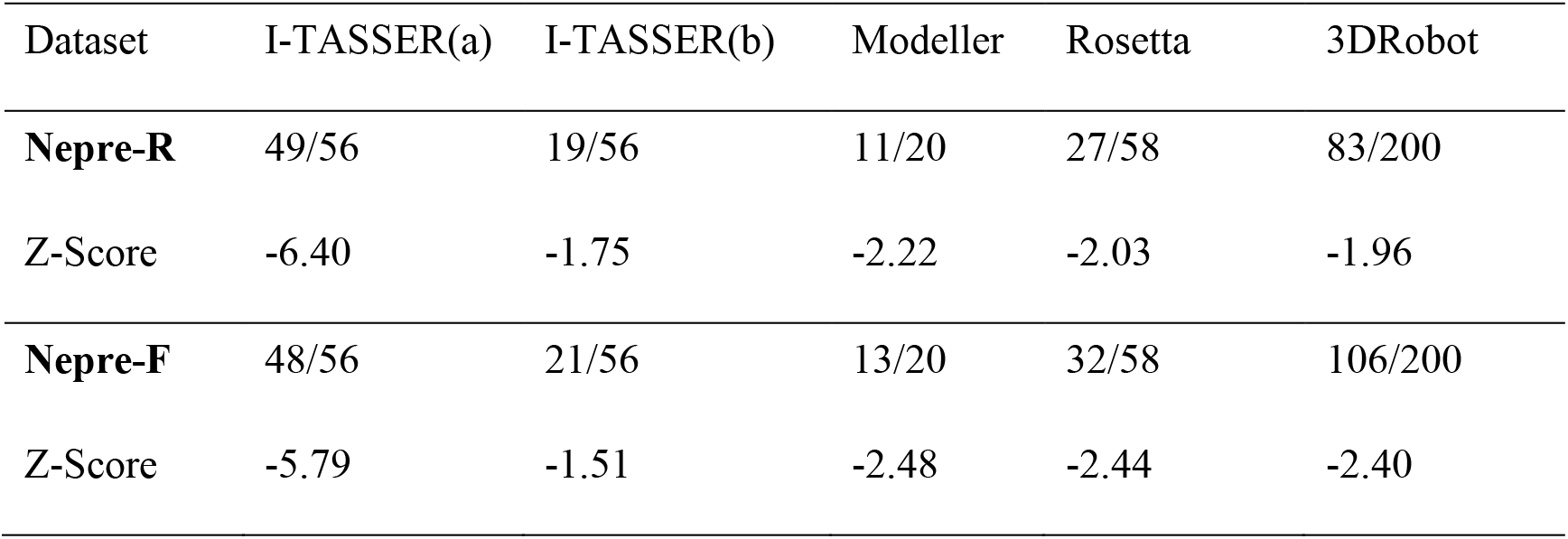
The comparison of performance of Nepre-R and Nepre-F.

As it is shown in Table 5, Nepre-R and Nepre-F have advantages in recognizing the native structures in decoy sets (represented as the number of TOP1 selected by each scoring function). Meanwhile, we also compared the sensitivity of the algorithm to check the ability of the algorithm to narrow down the native structure in a smaller ensemble selected based on the energy values. The smaller ensembles are composed of 1, 5, or 10 structure(s) with lowest energies. We compare the sensitivity of DFIRE, DOPE, RW, RWplus, Nepre (−R and −F), the results are summarized in Table 5 (TOP5 and TOP10, in addition to TOP1). Two phenomena were observed: (1) the Nepre algorithm performs well in all decoy sets, with Nepre-F showing slightly better results; (2) the success rate in selecting the native structure is increased as the ensemble size is increased. For example, when the ensemble is composed of 10 structures with lowest energies, the success cases is increased from 83 to 135 for 3DRobot decoy set using Nepre-R, and from 106 to 151 using Nepre-F for the same dataset. The same trend was observed for other algorithms as well.

**Table 5.** Performance comparison of different potentials.

|  |  | DFIRE | DOPE | RW | RWplus | Nepre-R | Nepre-F |
| --- | --- | --- | --- | --- | --- | --- | --- |
| I-TASSER(a) | TOP1 | 53/56 | 48/56 | 54/56 | 56/56 | 49/56 | 48/56 |
|  | TOP5 | 55/56 | 48/56 | 55/56 | 56/56 | 51/56 | 48/56 |
|  | TOP10 | 55/56 | 49/56 | 55/56 | 56/56 | 53/56 | 49/56 |
| I-TASSER(b) | TOP1 | 0/56 | 11/56 | 0/56 | 0/56 | 19/56 | 21/56 |
|  | TOP5 | 2/56 | 25/56 | 1/56 | 2/56 | 27/56 | 29/56 |
|  | TOP10 | 2/56 | 29/56 | 4/56 | 4/56 | 30/56 | 34/56 |
| Rosetta | TOP1 | 0/58 | 7/58 | 0/58 | 0/58 | 27/58 | 32/58 |
|  | TOP5 | 2/58 | 31/58 | 2/58 | 2/58 | 43/58 | 49/58 |
|  | TOP10 | 5/58 | 43/58 | 8/58 | 8/58 | 52/58 | 53/58 |
| Modeller | TOP1 | 2/20 | 6/20 | 2/20 | 2/20 | 11/20 | 13/20 |
|  | TOP5 | 4/20 | 8/20 | 4/20 | 4/20 | 16/20 | 18/20 |
|  | TOP10 | 6/20 | 10/20 | 6/20 | 7/20 | 17/20 | 18/20 |
| 3DRobot | TOP1 | 38/200 | 63/200 | 0/200 | 0/200 | 83/200 | 106/200 |
|  | TOP5 | 49/200 | 141/200 | 5/200 | 8/200 | 114/200 | 135/200 |
|  | TOP10 | 60/200 | 165/200 | 9/200 | 10/200 | 135/200 | 151/200 |

### Performance on CASP12 decoy datasets

Ten structures with resolution better than 2.5 Å were selected from CASP12 experiments. The decoys associated to these structures are the prediction models submitted by the participating teams in CASP12. The results of Nepre analysis on this dataset are summarized in Table 6. Two implementations, Nepre-R and Nepre-F, were both tested. In general, the Nepre with the universal fixed cutoff=6.0Å gave better results, succeeded in selecting native structures for five decoy sets (indicated by the RMSD=0Å cases in the last column of table 6). In the case of Nepre-R, the native structures were selected in only two out of ten decoy sets. Although this looks less promising, we found that the selected structures, which have the lowest energy in each decoy set, are very close to the corresponding native structures. In seven decoy sets, the selected structures have RMSD values within 3Å of the native structures. Similar results were found for the Nepre-F analysis. This testing indicates that the Nepre algorithm is generally applicable to the structure predictions.

**Table 6.** Performance of Nepre algorithm on 10 decoy sets in CASP12.

|  |  | Nepre-R |  |  | Nepre-F |  |  |
| --- | --- | --- | --- | --- | --- | --- | --- |
| No. | Decoy ID | Energy<br>for native<br>structure | Lowest<br>energy | RMSD* | Energy<br>for native<br>structure | Lowest<br>energy | RMSD* |
| 1 | T0860 | -42.34 | -43.48 | 1.52 | **-43.46** | **-43.46** | **0** |
| 2 | T0864 | -43.25 | -61.58 | 21.87 | -55.43 | -66.59 | 18.41 |
| 3 | T0869 | **-71.49** | **-71.49** | **0** | **-63.75** | **-63.75** | **0** |
| 4 | T0872 | -28.27 | -37.49 | 1.76 | -33.76 | -35.13 | 1.76 |
| 5 | T0877 | -41.06 | -45.43 | 2.77 | -38.02 | -46.36 | 3.11 |
| 6 | T0878 | -86.51 | -100.33 | 35.66 | **-119.68** | **-119.68** | **0** |
| 7 | T0879 | -72.81 | -78.95 | 1.39 | **-79.43** | **-79.43** | **0** |
| 8 | T0882 | -27.18 | -37.87 | 1.64 | -19.59 | -27.96 | 1.90 |
| 9 | T0900 | -27.98 | -34.56 | 13.63 | -32.81 | -35.03 | 7.99 |
| 10 | T0912 | **-126.92** | **-126.92** | **0** | **-171.49** | **-171.49** | **0** |

\* the RMSD is calculated between native and selected structures that have the lowest energies. The successful cases in selecting the native structure in each decoy set have an RMSD=0Å, and they are highlighted in bold font.

## DISCUSSIONS AND CONLCUSION

In protein structures, amino acids exhibit preferences on their neighboring amino acids, in both the types and the relative positioning of the amino acids. This property was systematically studied from the structures determined using experimental methods. Established on the results of neighbourhood preference, we have developed an new algorithm, Nepre, which is shown to be generally applicable in structure assessment. We have tested this algorithms using five published decoy sets and a new decoy set composed of 10 proteins with predicted models in CASP12. The excellent performance of Nepre algorithm has shown its potentials in structure predictions. The execution time is within 3-4 seconds for proteins in the tested decoy sets, including the PDB file parsing. Therefore, it is feasible to integrate the Nepre algorithm in model generation programs to sample the desired structure ensemble.

The Nepre algorithm was implemented in two forms, depending on the cutoff values that defines the neighborhood. The testing results have shown that the cutoff=6Å is an optimal choice for all 20 amino acids, regardless of the amino acid types. This was proved to be true in the case of the CASP12 decoy sets, which were not used in the determination of the optimal distance cutoff values. Surprisingly, the performance is even slightly better than the more sophisticated case of Nepre-R, which has distance cutoff values depending the neighboring amino acid sizes. Intuitively, the usage of type dependent cutoff values should gain more precious interactions between the amino acids, and therefore should lead to better performance of the algorithm. While we are still uncertain about the cause for inferior performance compared to its peer Nepre-F (with cutoff=6Å), there are several possible explanations. The radius values for each amino acid were obtained from the statistics in the protein structures, and the average values may not reflect the neighboring relation with other amino acids. For example, cysteine and serine each has two peak values, and using a single average value will result in some misrepresentation of the neighborhood (see Supplementary materials). The neighborhood described in this work is at residue level, since the distance is measured using center-to-center distance. Using amino acid type specific distance cutoff will enhance this residue level feature. On the other hand, using a universal fixed cutoff may reduce this strong selection, and the neighborhood is more uniformly defined. There is a necessity to carry on a detailed analysis to resolved this question.

The universal cutoff distance for the Nepre-F program was optimized by examing the performance of the program in five published decoy datasets. Besides this, the Nepre algorithm was not fine tuned in any other way. The distance dependence was considered during the neighborhood definition. It is reasonable to claim the Nepre algorithm is mainly depend on the orientation of the neighboring amino acids. The good performance of the algorithm indicate that the orientation is more critical for amino acids packed in a protein structure.

In summary, the neighborhood of amino acids in protein structures were statistically analyzed, and the discovered preferences were quantified using the neighbhoring amino acid types and relative positions. The neighborhood preference was then used to assess the structure quality for proteins, using a program implemented as Nepre. The Nepre programs showed excellent performance in selecting the native (or near native) structure from structure decoy sets. The algorithm can be generally applied in protein structure quality assessment and protein structure prediction studies. The source codes for Nepre is available via https://github.com/LiuLab-CSRC/ or upon request to the authors.

## Acknowledgements

This project is supported by National Natural Science Foundation of China (*Nos*. 11575021, U1530401, U1430237).

## Supplementary Information

**Figure S1.**
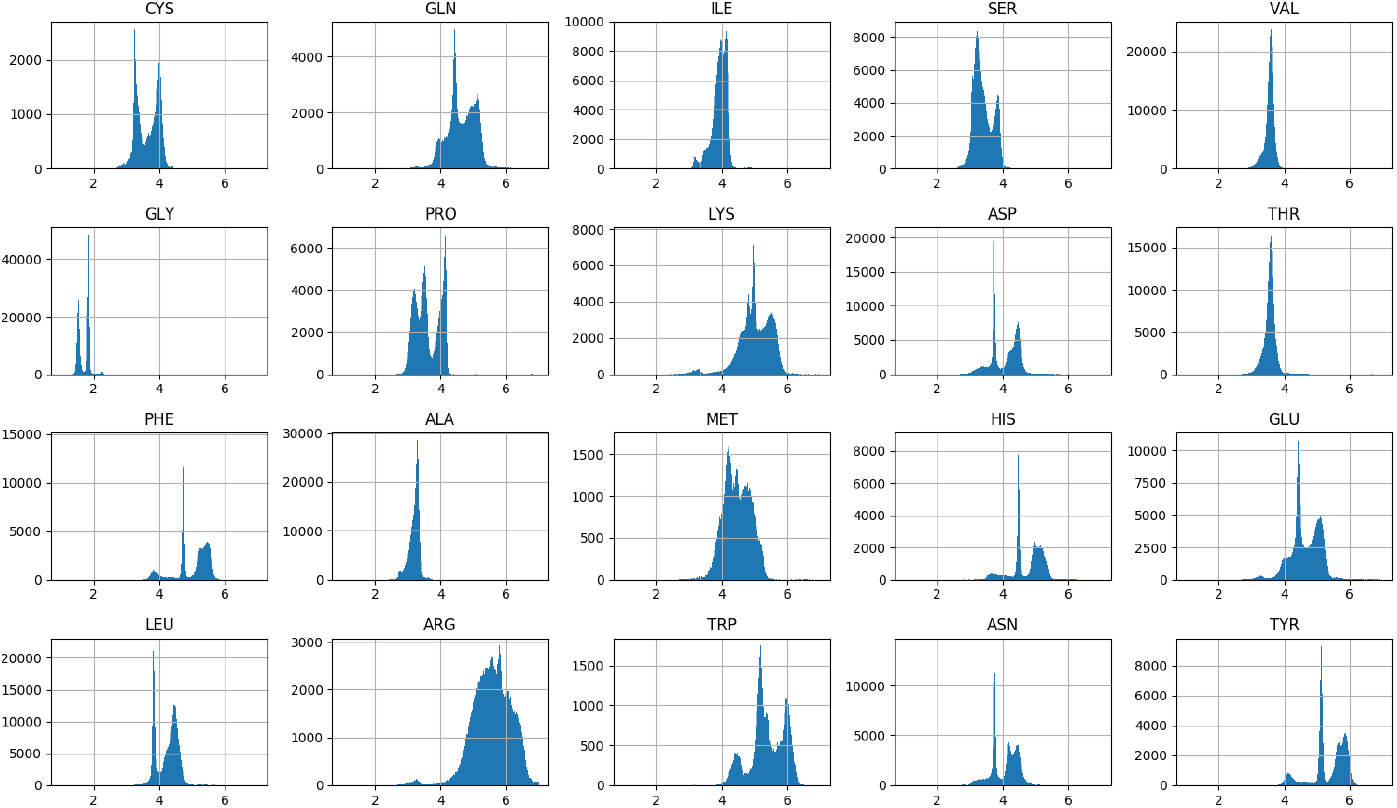
Distribution of amino acid radius. The statistics is based on the dataset composed of 30,000 high-resolution protein structures. The radius is defined as the largest distance between any atom and the geometry center of the amino acid.

**Figure S2.**
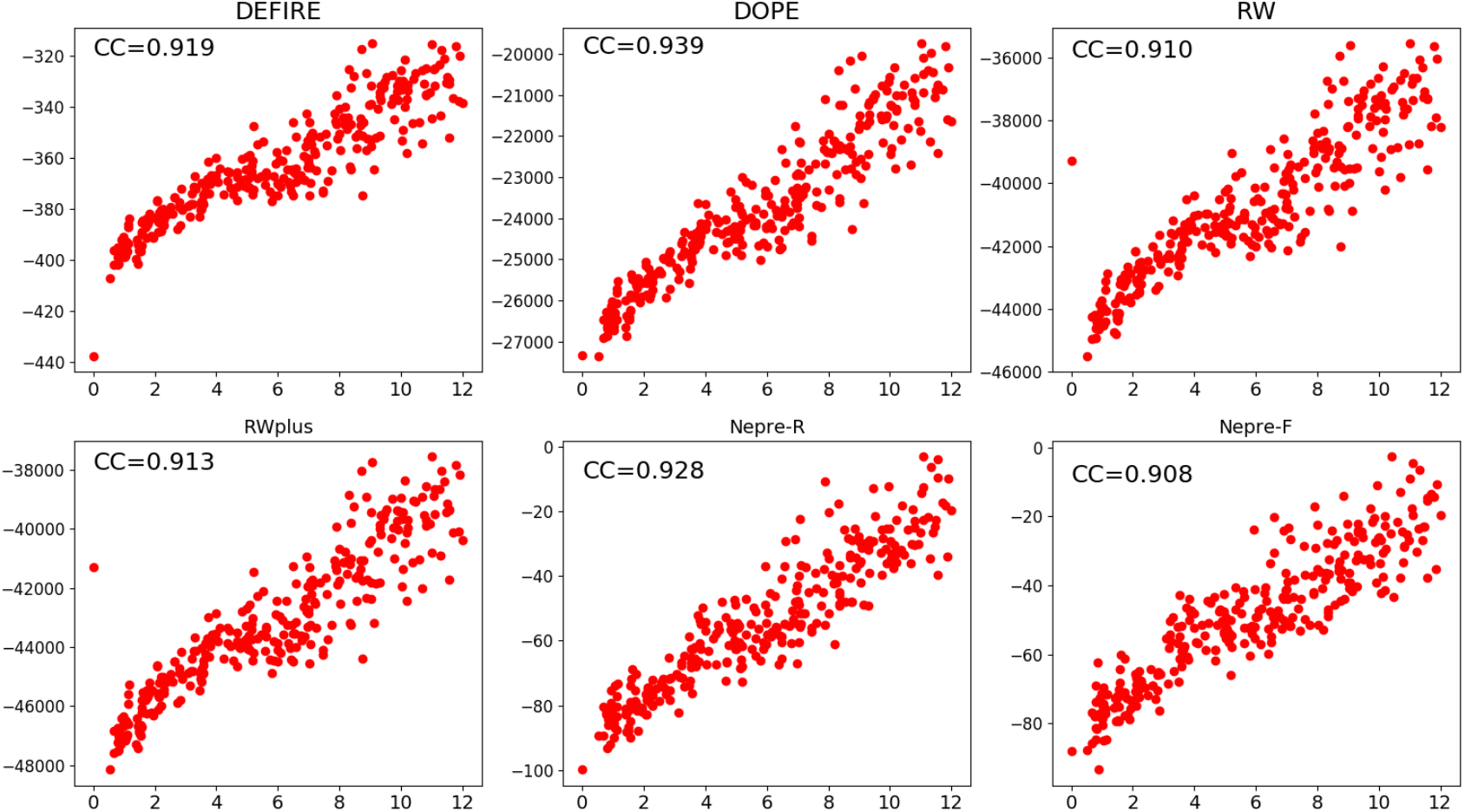
The correlation between scoring function and the structure difference compared to native state (measured using RMSD) for decoy set 1BYIA.

**Table S1.** Pearson coefficient on different datasets using different potentials.

|  | 3DRobot | I-TASSER(b) | ROSETTA | Modeller |
| --- | --- | --- | --- | --- |
| DFIRE | 0.810 | 0.754 | 0.769 | 0.776 |
| DOPE | 0.828 | 0.763 | 0.781 | 0.811 |
| RW | 0.802 | 0.756 | 0.771 | 0.781 |
| RWplus | 0.801 | 0.756 | 0.769 | 0.783 |
| Nepre-R | 0.767 | 0.647 | 0.691 | 0.769 |
| Nepre-F* | 0.734 | 0.609 | 0.671 | 0.730 |

\* Nepre-F used cutoff=6Å in this statistics.

**Table S2.** The native structure and Nepre selected decoy information for the 10 datasets from CASP12.

| No. | Structure | Native | Nepre-R<br>Minimum | Nepre-F<br>Minimum |
| --- | --- | --- | --- | --- |
| 1 | T0860 | 5fjl.pdb | T0860TS313_3 | T0860TS313_2 |
| 2 | T0864 | 5d9g.pdb | T0864TS384_1 | T0864TS450_2 |
| 3 | T0869 | 5j4a.pdb | T0869TS446_4 | T0869TS363_4 |
| 4 | T0872 | 5jmb.pdb | T0872TS384_5 | T0872TS384_5 |
| 5 | T0877 | 5nsj.pdb | T0877TS467_1 | T0877TS251_3 |
| 6 | T0878 | 5unb.pdb | T0878TS450_5 | T0878TS247_3 |
| 7 | T0879 | 5jmu.pdb | T0879TS005_2 | T0879TS005_5 |
| 8 | T0882 | 5g3q.pdb | T0882TS243_1 | T0882TS247_5 |
| 9 | T0900 | 5aot.pdb | T0900TS282_3 | T0900TS247_5 |
| 10 | T0912 | 5mqp.pdb | T0912TS475_4 | T0912TS411_1 |

